# Testing hypotheses on population dynamics: A test for the inhomogeneous Poisson point process model

**DOI:** 10.1101/234948

**Authors:** Niklas Hohmann

## Abstract

In this paper, a test for hypotheses on population dynamics is presented alongside an implementation of said test for R. The test is based on the assumption that the sample, consisting of points on a time axis, is a realization of a Poisson point process (PPP). There are no restrictions on the shapes of the rate functions that are regulating the PPP, type 2 errors can be calculated and the test is optimal in the sense that it is a uniform most powerful (UMP) test. So for every significance level α, the presented test has a lower type 2 error than every other test having the same significance level α. The test is applicable to all models based on PPPs, including models in spatial dimensions. It can be generalized and expanded in different ways, such as testing larger hypotheses, incorporating prior knowledge, and constructing confidence regions that can be used to obtain upper or lower bounds on rate functions.

## 1 Introduction

**Note:** *for clearer language, the explanations in this paper will be restricted to the case where samples consist of single points on the time axis. The case for more general models like models in spatial dimensions follows mutatis mutandis*. Imagine an observation point at which the occurence of individuals from a certain species throughout time is documented. The actual observations times are random, but depend strongly on the total number of individuals of that species around the observation point. By applying a limiting theorem on a simple sampling procedure, it can be shown that the stochastic pattern underlying the observation times is a Poisson point process (PPP) with rate λ. [Meester(2008), ch. 7.1] The parameter λ represents the average number of individuals observed per time unit and can be estimated with ordinary statistical methods for the Poisson or the exponential distribution.

Now, assume the number of individuals around the observation point is changing over time, for instance because the species has a migratory habit. Then the number of observed individuals will change over time, since the number of individuals around the observation point changes over time. By arguments similar to those above, it is plausible that the stochastic pattern underlying the observation times is an inhomogeneous PPP. The changing observation rates of this process are formalized by a rate function *f* (*t*), where *t* is the time. For this PPP, the probability of observing a certain number of individuals in a interval [*t*_0_, *t*_1_] is given by the Poisson distribution with parameter 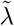, [Klenke(2008), def. 24.10]^1^ which is defined as

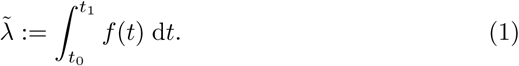

In the case of changing observation rates, knowing the rate function of the underlying PPP allows important conclusions to be drawn. For example, knowing the migratory habit of the observed species is equivalent to knowing the rate function of the underlying PPP.

One approach to retrieve information about the unknown rate functions is to estimate them. But based on the choice of the estimation method used, this either leads to estimates that are not unique, trivial estimates^2^ or to the usage of models based on strong a priori assumptions^3^ that are usually hard to justify. To avoid those problems, the focus is shifted from estimating a rate function to testing competing rate functions. By doing this, the questions which arise from using different estimation methods are avoided in favor of the question which predefined rate function is more likely to have generated an observed sample.^4^.

## 2 The test

### 2.1 Description

Assume there are two hypotheses *f*_0_,*f*_1_ about the unknown rate function *f* underlying a PPP. The function *f*_0_ will be treated as the null hypothesis and the function *f*_1_ as the alternative. This leads to the test of

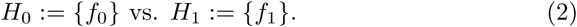

Let t = *t*_1_,*t*_2_,…*t_N_* be a sample containing the observation times of the individuals from the species of interest. Define

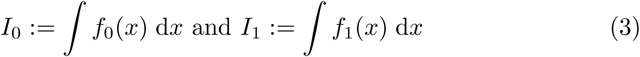

and

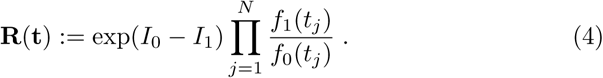

Then the test *φ* is given by

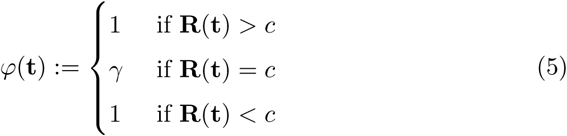

with *c* and *γ* chosen in a way that the desired significance level *α* is met. In the further explanations, this test will be called p3r2 test as a abbreviation of **P**oisson **p**oint **p**rocess **r**ate **r**atio test.

### 2.2 Properties and implementation

The p3r2 test is a uniform most powerful (UMP) test for the model of a PPP with two competing rate functions *f*_0_ and *f*_1_. So for any significance level *α*, the p3r2 test has a smaller type 2 error than any other test with significance level *α*. This follows from equation (13), which makes the test a likelihood-ratio test in a binary model and therefore UMP (Neyman-Pearson lemma, see [Liese and Miescke(2008), theorem 2.45]).

According to Stein’s theorem ( [Liese and Miescke(2008), theorem 8.75]), the type 2 error converges exponentially fast to zero:

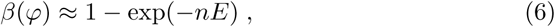

with *E* being the Kullback-Leibler divergence and *n* the number of samples. The value of *E* is

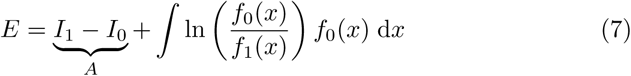

and corresponds to how well *f*_0_ and *f*_1_ can be distinguished. Note that the first part of that equation (marked with A) is solely dependent on the integral over *f*_0_ and *f*_1_, which is generating the overall number of observations, and not of the shape of *f*_0_ or *f*_1_. So one part of how well the functions can be distinguished is dependent on their ability to generate a plausible number of observations, and not information about where those observations are made. The part marked with *A* in equation (7) can therefore be called the sample size share of the Kullback-Leibler divergence. So if only the shapes of two functions should be tested, they should be chosen so that *A* does vanish. To make this easier, a rescaling option in the implementation is available.^5^ The effects of sample size and resacling are explained and discussed in the example in the section below. The p3r2 test and a modified version for binned data is implemented for R and can be found in the appendix. The central part of the implementation is the simulation of the distribution of **R**, given *f_i_* for *i* ∈ {0,1}. For this, a PPP with rate function *f_i_* is simulated, using a method similar to inverse tranform sampling.^6^ Then **R** is evaluated at the simulated sample. This is repeated a large number of times. Then the empirical cumulative distribution function of all those values will converge to the cumulative distribution function of **R**, given *f_i_*. Convergence follows from the Glivenko-Cantelli theorem (see [Klenke(2008), theorem 5.23]), convergence rates can be derived from the Dvoretzky-Kiefer-Wolfowitz inequality (see [Kosorok(2008), theorem 11.6]). For more details on the simulation of PPPs, numerical details and considerations see the documentation of the code as well as comments in the code.

## 3 Example

As an example, the test is applied to a dataset from [Solow(1993)], which consists of sighting data of the black-footed ferret. The data was rescaled to the time unit of one year with the first year of observation being the interval between zero and one. The observation period ended after thirteen years. The test is applied in two slightly different scenarios to emphasize the differences in model assumptions and their influence on the results of the test.

In the first scenario, the rate function of the null hypothesis is chosen to be constant over the whole observation period. Since there are four sightings in the first year, the value of the rate function is set to four. So

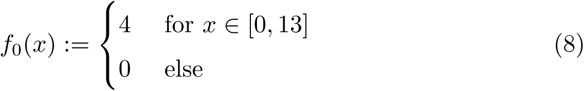

The rate function of the alternative is chosen to be constant for the first year. After that, it is declining until it vanishes at *t* = 13. Again, the sighting rate in the first year is set to four. So

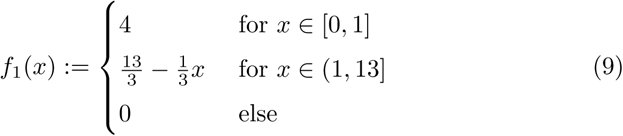

The rate functions are displayed in figure 1. Applying the test^7^ returns a p-value of *p* = 0.00035, the type 2 error at level *α* = 0.05 is 0.00001. The value of the test statistic evaluated with the sample is approximately 39 and the critical value for the significance level *α* = 0.05 is *c* = 0.00015. This is strong evidence to reject the null hypothesis. Plots of the distribution functions and type 1 versus type 2 error are shown in figure 2 and figure 3.

**Figure 1:**
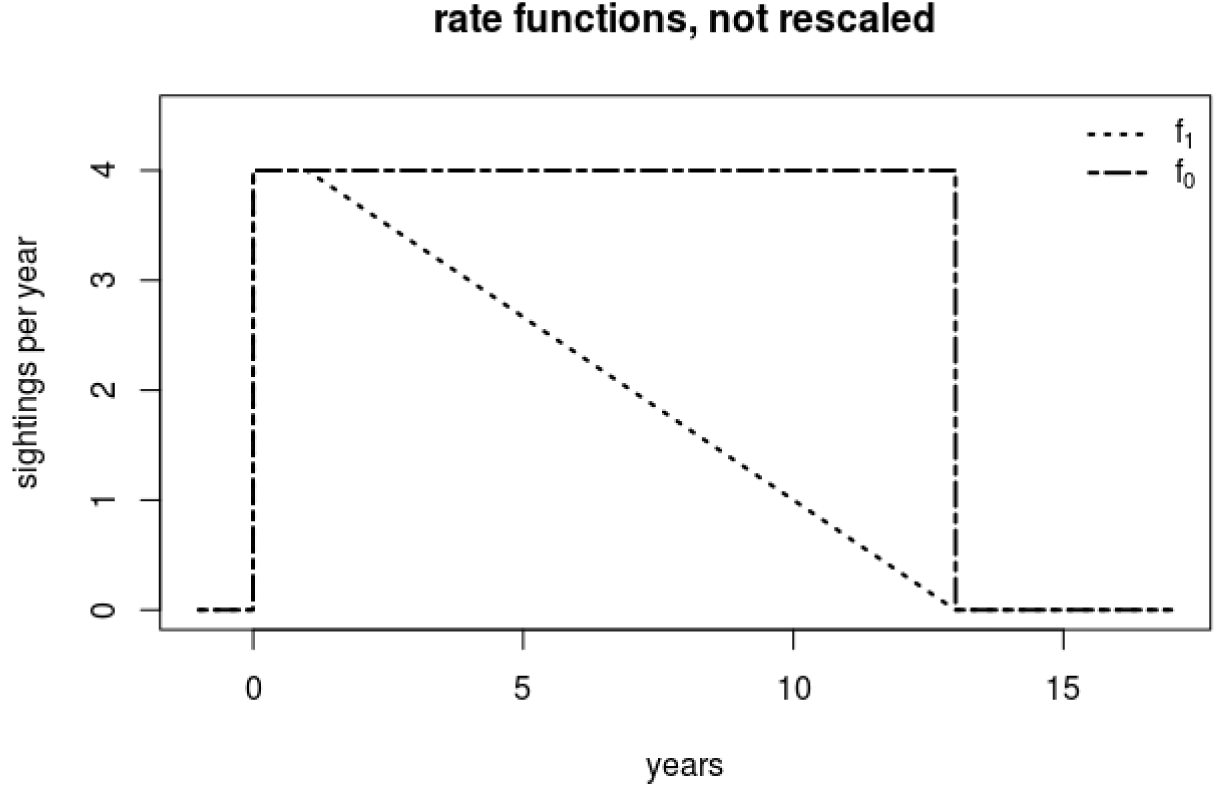
The unchanged rate functions from the first scenario.

**Figure 2:**
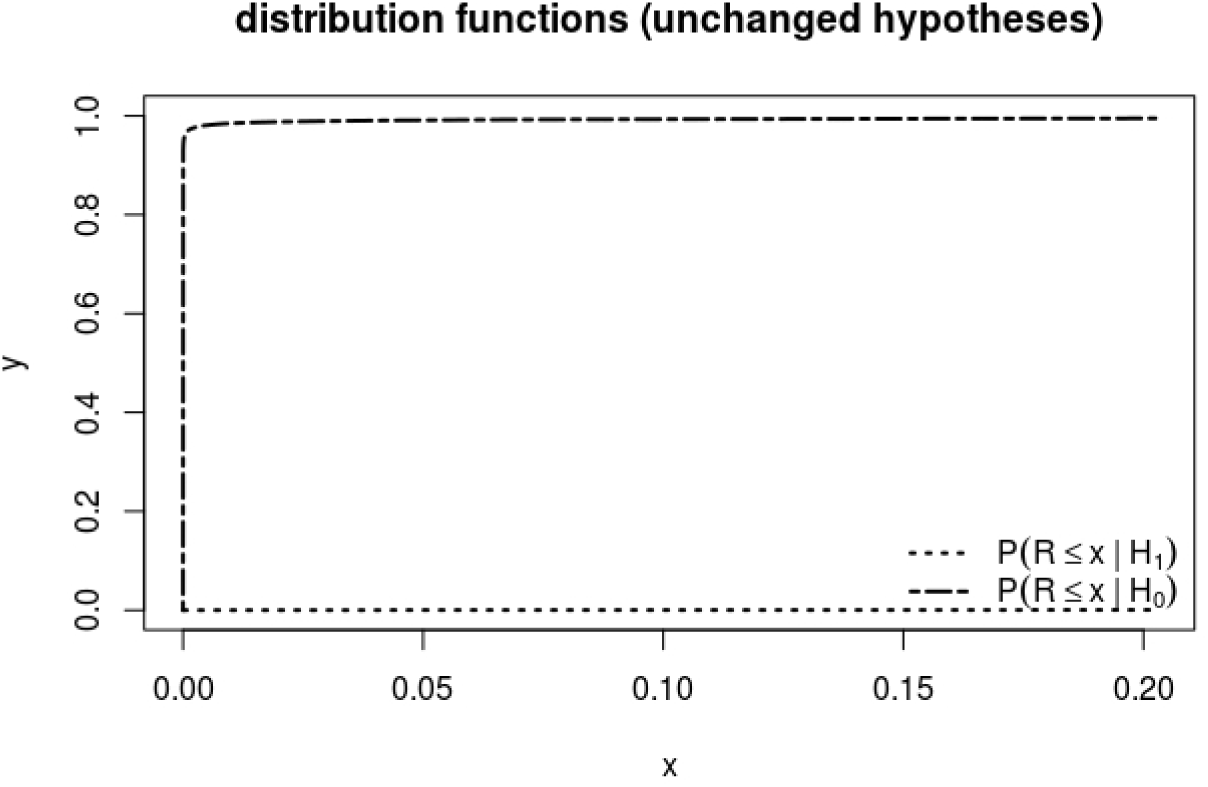
Distribution functions of the test statistic **R** for the unchanged rate functions from the first scenario.

**Figure 3:**
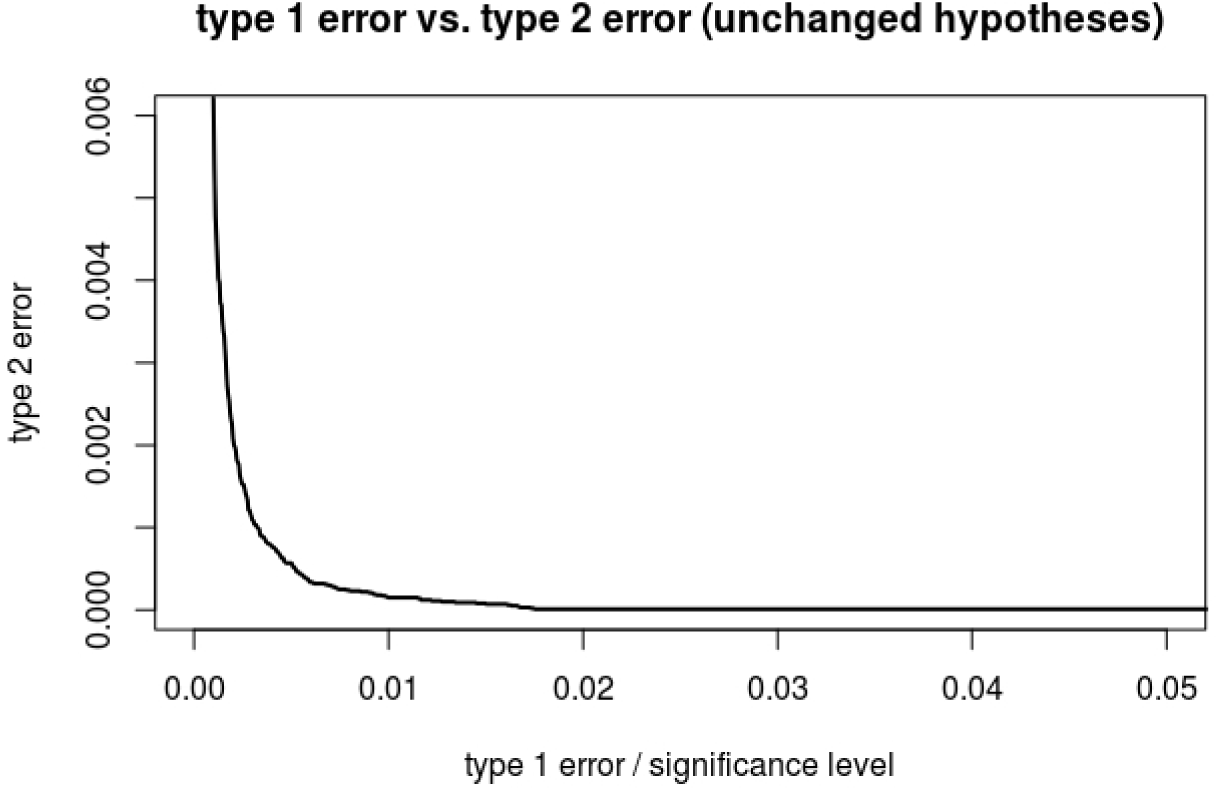
Plot of type 1 error vs. type 2 error for the test with the unchanged rate functions from the first scenario.

With the hypotheses chosen as described above, equation (3) returns *I*_0_ = 52 and *I*_1_ = 28. From equation (1) and the properties of the Poisson distribution it can be derived that the average number of sightings per sample under the null hypothesis is 52, whereas the average number of sightings per sample under the alternative is 28. In the data from Solow, there were 28 sightings, which is exactly the average number of sightings under the alternative. Assuming that the null hypothesis is true, it would be a very unlikely event to have that few sightings. In fact, the probability for a sample with less than 28 or more than 76 sightings (52 ± 24) is lower than 0.0009 given that the null hypothesis is true.

So the chances of rejecting the null hypothesis are amplified by the choice of a null hypothesis which is implausible in terms of average sample size.^8^

To study this effect, the test is applied a second time with modified rate functions. Now, they are rescaled so that the average number of sightings is the observed number of sightings. This is done by using the rescaling option which was discussed in the follow-up of equation (7) and is integrated into the implementation of the test. There is no effect of rescaling on *f*_1_, since *I*_1_ already matches the number of sightings in the data set. The constant function *f*_0_ is rescaled such that *I*_0_ = 28, so *f*_0_ has got a constant rate of 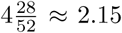. The rescaled rate functions are shwon in figure 4. Running the test again with the rescaled hypotheses,^9^ the p-value is *p* = 0.1494 with a type 2 error of 0.0249 at a significance level of *α* = 0.05. The value of the test statistic evaluated with the sample is approximately 0.056 and the critical value for the significance level *α* = 0.05 is *c* = 0.73. Plots of the distribution functions and type 1 versus type 2 error are shown in figure 5 and figure 6. This time, the null hypothesis can not be rejected.

**Figure 4:**
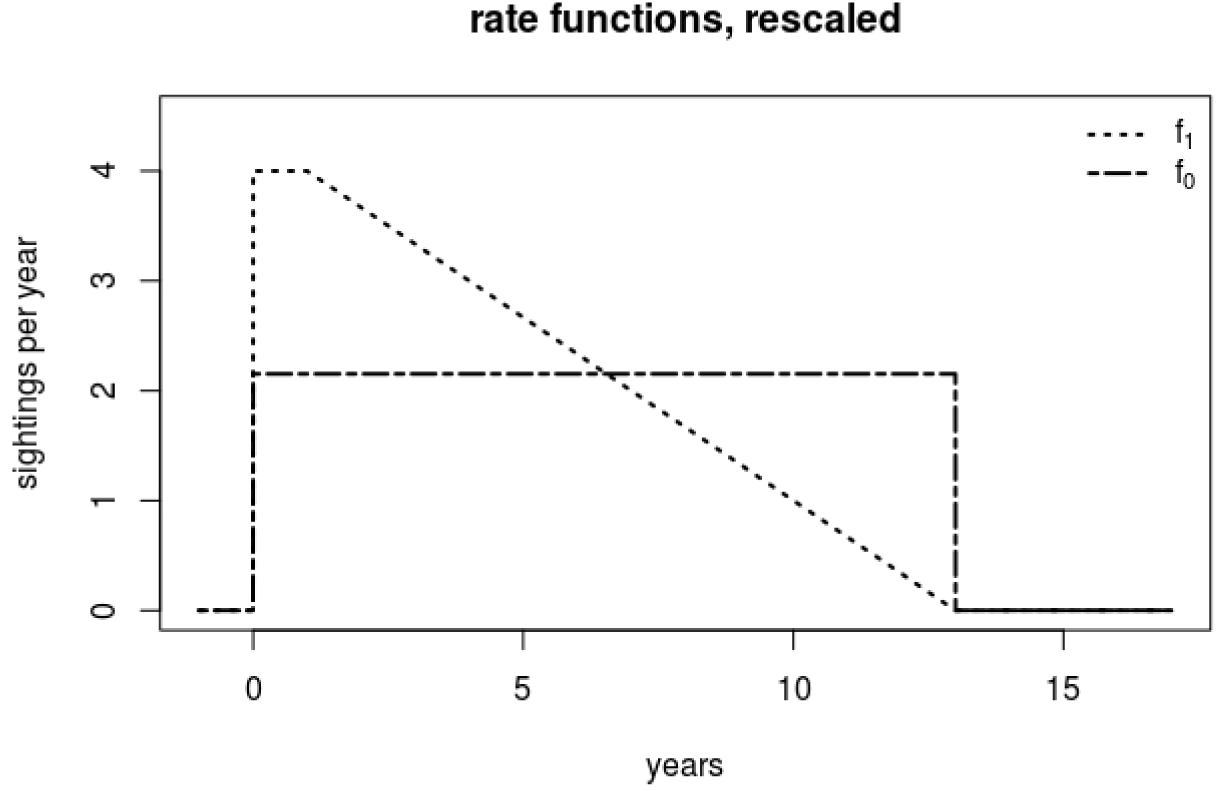
The rescaled rate functions.

**Figure 5:**
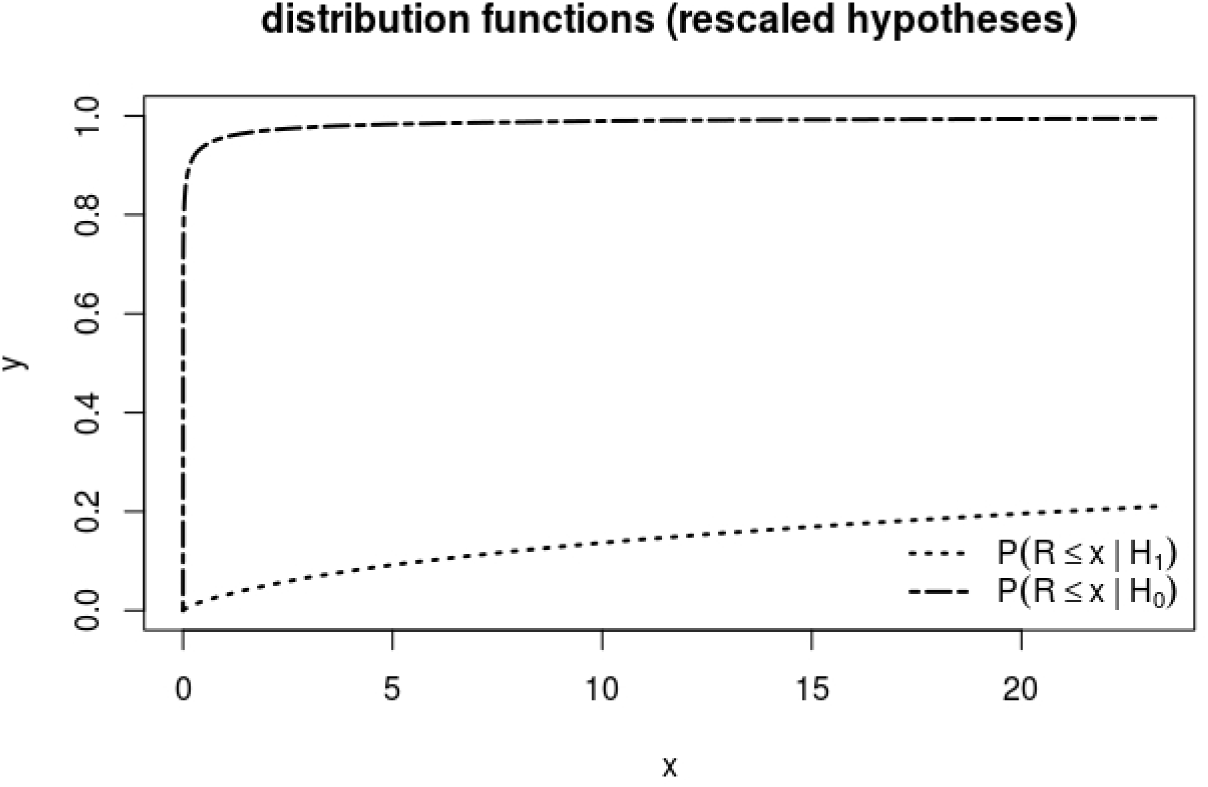
Distribution functions of the test statistic **R** for the rescaled rate functions.

**Figure 6:**
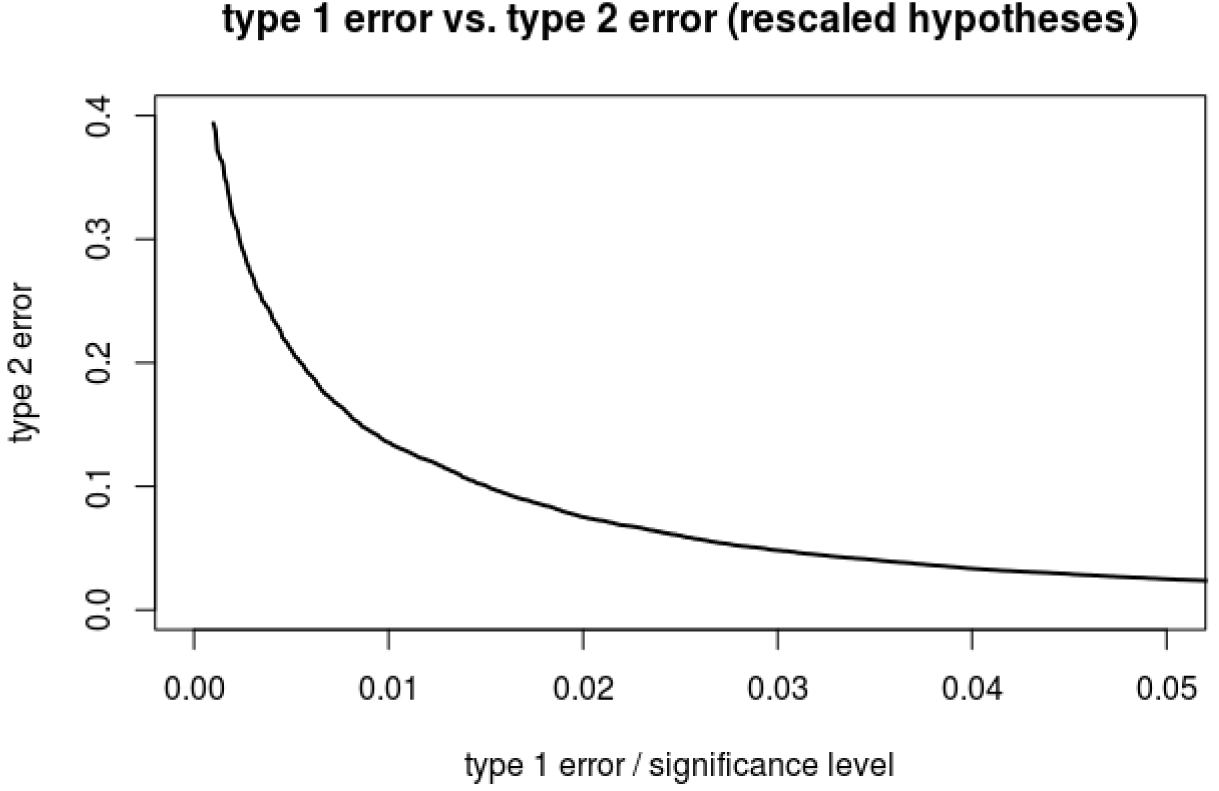
Plot of type 1 error vs. type 2 error for the test with the rescaled rate functions. Note the change in scale on the y axis in comparison to figure 3.

This example illustrates the importance of properly choosing hypotheses, especially since the important factors *I*_0_ and *I*_1_ are easily misestimated.^10^ When there are no explicit reasons to choose hypotheses that generate different average sighting numbers, rescaling should always be used since it focuses the testing problem more on the shape of the hypotheses than on the overall number of sightings. Note that rescaling makes the testing problem harder, since it reduces the Kullback-Leibler distance (eliminating *A* in equation (7)). This makes the null hypothesis and the alternative harder to distinguish. Rescaling basically reduces the amount of information available to differentiate between the hypotheses since it makes it impossble to draw information from sample size.

## 4 Sketch of mathematical structures

This section briefly sketches the used structures. For a more detailed discussion see the pdf file *math* in the appendix. For *i* ∈ {0,1}, define the measures

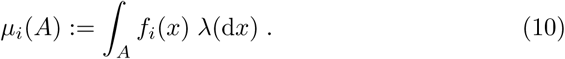

Here, λ is the Lebesgue measure. These measures serve as intensity measures for PPPs that are formalized as random measures *X_i_*.^11^ The *X_i_* are measure valued random elements and therefore posess distributions *P_i_* on the set of measures. It can be shown that there is a probability distribution *P_M_* on the set of all measures satisfying *P_i_* ≪ *P_M_*, so the Radon-Nikodym theorem guarantees the existence of a density function *d_i_*. It is given by

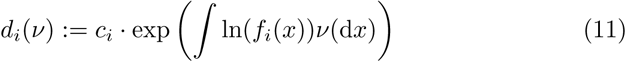

with

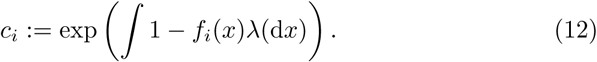

With this, the likelihood-ratio function can be defined:

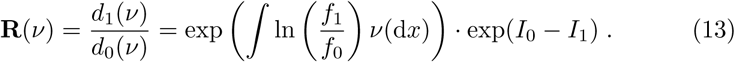

Since all relevant measures *v* are sums of Dirac measures, this reduces to the representation of the test statistic in (4). Therefore the test as defined in equation 5 is a likelihood-ratio test with simple hypotheses.

## 5 Possible extensions

In this section, possible extensions of the p3r2 test and of the structures that were used to construct it are discussed.

### 5.1 Larger hypotheses

The p3r2 test for simple hypotheses can be expanded in the sense that larger sets of rate functions are feasible as hypotheses. The null hypothesis can be expanded from one function *f*_0_ to

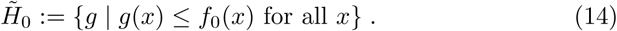

The p3r2 test is still a UMP test if it is used to test 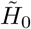 against *H*_1_ (as defined in equation (2)). This is because for every 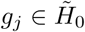

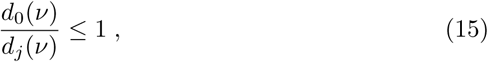

so the optimality for larger null hypotheses can be derived from a monotonicity argument.

Expanding the alternative to

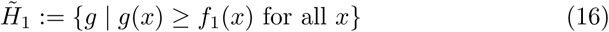

is also possible, but it is not clear if the p3r2 test is still optimal in any sense when it is applied in this situation. One way to establish maximin optimality for the test in this general situation might be to use concepts from robust testing. By looking at 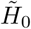 and 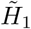 as neighbourhoods of some not further specified probability distributions, the test of *f*_0_ against *f*_1_ might be a the worst-case scenario for those neighbourhoods and therefore maximin for testing 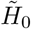 vs. 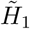.

### 5.2 Incorporating partial knowledge

Partial knowledge of relevant processes can also be incorporated into the test. For this, the rate function 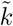 of the known process is combined with the assumed rate functions 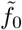 and 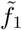 to create the rate functions *f*_0_ and *f*_1_ that are being tested. The test results for *f*_0_/*f*_1_ are then equal to the test results for 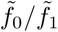. It is important to know how the known and the unknown process interact, since this determines the way that 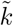 and 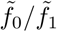 are combined to form *f*_0_/*f*_1_. In most cases, the combination will be a simple multiplication of the functions, since this corresponds to a multi stage sampling. This combination of known processes and rate functions can of course be varied in multiple ways. Examples might be multiple known influences overlying each other or different known processes for the null hypothesis and the alternative. A basic example is presented in the pdf file *docp3r2contR* which can be found in the appendix.

### 5.3 Exponential family

The density as defined in equation (11) can be interpreted as a ∞-parametric exponential family. With the abbreviations

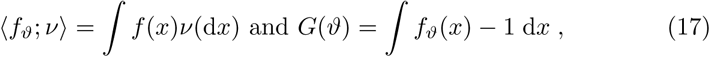

it can be represented as

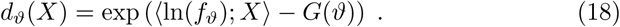

This is equivalent to the representation of a member of the exponential family as shown in [Liese and Miescke(2008), p. 3 eq. 1.6].

This additional structure can for example be exploited by reducing the dimensionality of the exponential family to obtain stronger results. One possibility is to fix a suitable *g* and define *f_ϑ_*:= *g^ϑ^* with *ϑ* > 0. Then

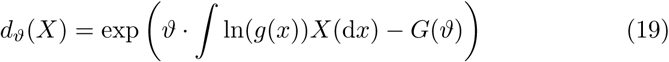

is a member of a oneparametric exponential family and has got a monotone likelihood-ratio in *∫* ln(*g*(*x*))*X*(*dx*). So all results for the oneparametric exponential family can be applied to this situation.

### 5.4 Confidence regions

Every confidence region is dual to a family of tests, see [Liese and Miescke(2008), p. 431-437]. With this duality, optimality results for confidence regions transform to optimality results for tests and vice versa.

Therefore the p3r2 test can be used to obtain optimal confidence regions for rate functions. Note that those confidence regions are not intervals or subsets of ℝ^*n*^, but sets of functions. So for a given set of rate functions 𝓕, the confidence region is a subset 𝓒 ⊂ 𝓕. One application for those confidence regions might be to obtain upper or lower bounds (with confidence level 1 — *α*) on the rate function that has generated a given sample.

## 6 Discussion

### 6.1 Common misconceptions

There are two common misconceptions about the interpretation of the test. The first misconception concerns the relation of the rate functions as a whole and their isolated properties. It is not possible to derive any conclusions about single properties of the rate functions from the test. They should be treated as indivisible objects. As an example, assume the population dynamics of a species is being tested. The null hypothesis *f*_0_ has been rejected and *f*_1_ is zero from *t* =10 an. That does not mean that the test returned the result that the species is extinct after ten time units. The test result concerns *f*_1_ as a whole. Conclusions about single properties, in this case extinction times, are not supported by the test result.

The second misconception concerns the interpretation of the rate functions. They are the the combination of all influences on the process. If some of these influences are unknown to the user or are being neglegted, the interpretation of the test result will be wrong. As an example, look at fossil occurences in geology. Here, the continuing absence of fossils is naively interpreted as the corresponding species being extinct. But there are a lot more influences on fossilisation than the number of individuals alive at that time. Some are changes in sedimentation and facies. So what is being tested here is the combination of all those influences that have generated the sample at which the test is being performed. One example of a misinterpretation can be found in the documentation for the test.^12^

### 6.2 Testing versus estimating

As pointed out in the introduction, testing has some advantages over estimating. In the Poisson point process model, estimation leads to a dilemma. On the one hand, using general estimation methods like martingale estimators or maximum-likelihood estimators leads to estimates that are not smooth and therefore unrealistic. On the other hand, methods aiming at smooth rate functions are often based on parametrisations and a priori assumptions, which leads to a loss of generality.

In contrast to that, testing allows the choice of smooth hypotheses without a loss of generality. But it does not give a absolute measure of quality. The measure of quality obtained from testing is a relative measure, since it just returns which of the chosen hypotheses is more plausible. However, this does not necessarily mean that any of the hypotheses that were tested is good in the sense that it is close to the real rate function. So in a broad sense, by changing from estimation to testing, the burden of quality is shifted from the choice of a proper method of estimation to the choice of proper hypotheses.

### 6.3 Summary

From a mathematical point of view, the presented test is the best possible test for a very flexible and plausible point process model that is underlying many processes in nature. In addition to that, the possibility to further exploit the structures developed show that the approach used still has great potential. The test and the implementation are almost unrestricted in terms of the choice of the rate functions. This allows the use of the test for a wide range of problems and in many different settings. Especially the incorporation of prior knowledge can be applied to different problems. One such problem might be the study of sampling bias or the incorporation of the known fossilisation bias in paleontology. This has been studied extensively by many authors [Wang and Marshall(2016)] [Wagner and Lyons(2011)] and remains a problem.

However, to avoid possible misuse of the test^13^ or misinterpretation of the results^14^ the test should always be used with care and thought. [Wagner and Lyons(2011)] wieso [Wang and Marshall(2016)]

## Footnotes

1 In combination with equation (10)

2 That includes estimates for the integrated rate function using martingale estimators

3 Such as parametrisations of a predefined set of rate functions as in [Solow(1993)]

4 For a short discussion of testing versus estimating see the discussion.

5 For more details, see the corresponding documentations of the R functions

6 A self-contained function for the simulation of a PPP with arbitrary rate function can also be found in the appendix

7 Parameters were *nor*= 100000, rescale FALSE, *type2* TRUE

8 This line of argument can be seen as a implicit maximum-likelihood heuristic for the choice of the hypothesis

9 Parameters were *nor*= 100000, rescale TRUE, *type2* TRUE

10 This may be due to the fact that it is harder to estimate area than length.

11 For a brief introduction into random measures see [Klenke(2008), chapter 24]; a very detailed discussion can be found in [Kallenberg(2017)]

12 See the file *docp3r2contR.pdf*

13 See the discussion of the rescale option in the example section

14 See discussion above

## 7 Appendix

The appendix contains the following files:

- an R file named *pppsample* for the simulation of a sample from a PPP with arbitrary rate function. The corresponding documentation can be found in the pdf file *docpppsampleR*
- an R file named *p3r2cont* containing the implementation of the test (in continuous time) as defined in (5). The corresponding documentation can be found in the pdf file *docp3r2contR*
- an R file named *p3r2bin* containing the implementation of the binned version of the test. The corresponding documentation can be found in the pdf file *docp3r2binR*
- an R file named *reevalp3r2* for performing multiple tests on unchanged hypotheses. The corresponding documentation can be found in the pdf file *docreevalp3r2R*
- a pdf file named *math* containing the mathematical proofs needed for the construction of the test

